# Transcription factor interactions explain the context-dependent activity of CRX binding sites

**DOI:** 10.1101/2023.03.05.531194

**Authors:** Kaiser J. Loell, Ryan Z. Friedman, Connie A. Myers, Joseph C. Corbo, Barak A. Cohen, Michael A. White

## Abstract

The effects of transcription factor binding sites (TFBSs) on the activity of a *cis*-regulatory element (CRE) depend on the local sequence context. In rod photoreceptors, binding sites for the transcription factor (TF) Cone-rod homeobox (CRX) occur in both enhancers and silencers, but the sequence context that determines whether CRX binding sites contribute to activation or repression of transcription is not understood. To investigate the context-dependent activity of CRX sites, we fit neural network-based models to the activities of synthetic CREs composed of photoreceptor TFBSs. The models revealed that CRX binding sites consistently make positive, independent contributions to CRE activity, while negative homotypic interactions between sites cause CREs composed of multiple CRX sites to function as silencers. The effects of negative homotypic interactions can be overcome by the presence of other TFBSs that either interact cooperatively with CRX sites or make independent positive contributions to activity. The context-dependent activity of CRX sites is thus determined by the balance between positive heterotypic interactions, independent contributions of TFBSs, and negative homotypic interactions. Our findings explain observed patterns of activity among genomic CRX-bound enhancers and silencers, and suggest that enhancers may require diverse TFBSs to overcome negative homotypic interactions between TFBSs.

## Introduction

A typical mammalian transcription factor (TF) binds hundreds or thousands of *cis*-regulatory elements (CREs) in the genome [1–3]. CREs that are bound by the same TF vary widely in their activity, and can include strong enhancers, transcriptional silencers, or sequences with weak or no *cis*-regulatory activity [3–20]. Such dramatic functional differences among CREs with similar TF binding sites (TFBSs) show that local sequence context modulates the contribution of a TFBS to *cis*-regulatory activity, yet how this occurs is not well understood. Proposed models of *cis*-regulatory grammar that may account for context-dependence vary in their emphasis on the importance of interactions between TFs, and they suggest different degrees of flexibility in the possible functional arrangements of TFBSs [21–24]. The enhanceosome model proposes that strict geometrical constraints determine whether CRE-bound TFs can activate transcription, suggesting that context-dependent effects of TFBSs are strongly influenced by highly specific interactions between them [21–23,25]. The contrasting billboard model proposes that active CREs are defined by the presence of a sufficient number of bound TFs, with no strong constraints governing their arrangement [24]. The billboard model implies that the context of a TFBS is determined primarily by additive effects of the surrounding TFBSs, with few specific interactions between sites. Other models of *cis*-regulatory grammar propose that individual TFBSs are weak on their own and depend on strong cooperative interactions [26], that particular TFBSs recruit specific, required transcriptional cofactors [11], or that the balance among sites for transcriptional activators and repressors determines whether a CREs is an enhancer or silencer [27–29]. The degree to which these proposed features of *cis*-regulatory grammar modulate the local context within a CRE is not well understood. As a result, accurately predicting the activity of CREs or the effects of genetic variants in TFBSs remains an unsolved problem.

Local sequence context has strong effects on the function of binding sites for the retinal TF Cone-rod homeobox (CRX) [30–32]. CRX is paired-type K50 homeodomain TF and a critical regulator of transcription in multiple retinal cell types, where it contributes to both activation and repression of cell type-specific genes [8,30,32–41]. Using massively parallel reporter assays (MPRAs), we previously found that genomic CRX-bound sequences include strong enhancers and silencers [13,18,42,43]. The activities of these CREs, whether activating or repressing, depend on both CRX binding sites and CRX protein, which demonstrates that the effects of CRX sites are modulated by context [13,42]. Yet how local sequence context determines whether a CRX-bound region functions as an enhancer or a silencer is poorly understood. We previously found that synthetic CREs with sites for CRX and the rod photoreceptor-specific leucine zipper TF NRL were often strongly activating [13], and CRX cooperatively interacts with NRL at some promoters [30,36,38,39,44–46]. We also found that CRX-bound silencers tend to contain more copies of the CRX motif than CRX-bound enhancers [13,18,42], while CRX-bound enhancers are enriched in sites for other TFs relative to silencers [18]. However, it is unclear why CRX binding sites have an activating effect at some elements and a repressing effect at others. We hypothesized that interactions between CRX and other co-bound TFs determine whether a sequence functions as an enhancer or silencer, and we sought to capture those interactions in a quantitative model.

A key advantage of synthetic CREs is that their binding site composition can be systematically varied to generate informative training data for models of *cis*-regulatory grammars. We used our MPRA data from synthetic CREs to train neural network-based models using MAVE-NN, a modeling framework designed to efficiently model data from massively parallel functional assays [47]. We find that the effects of CRX sites are explained by a model that includes positive, additive contributions of individual TFBSs, negative homotypic interactions between sites for the same TFs, and positive heterotypic interactions between sites for different TFs. The model explains the observations that CRX sites produce context-dependent activation and repression, and that the addition of an NRL site converts silencers to enhancers. The model also accounts for our finding that CRX-bound enhancers have sites for diverse TFBS, while CRX-bound silencers lack this diversity. More generally, our results suggest that context-dependent activity of binding sites for transcriptional activators can be explained by the balance between the negative effects of interactions between sites for the same TF, the positive effects of individual TFBSs, and heterotypic cooperativity between sites.

## Results

### Positive heterotypic and negative homotypic interactions explain the effects of CRX and NRL sites on expression

We previously reported that both genomic and synthetic CREs with many binding sites for CRX tend to act as silencers, while CREs with fewer CRX sites tend to act as enhancers [13,18]. Our prior results from a reporter library of 1,299 synthetic CREs showed that sequences composed of only CRX and NRL binding sites exhibit activity that ranges from strong activation to repression [13]. These CREs were tested by MPRA in mouse retinal explants, which preserve all retinal cell types and cell type-specific TFs that comprise the native context in which CRX is active. We observed that sequential addition of CRX sites upstream of a basal promoter led first to increased activation and then to repression below basal levels when three or four CRX sites were present (**Fig. 1A and B**). Repressive CREs with four CRX sites could be converted to strongly activating sequences by replacing one CRX site with a site for NRL. Synthetic sequences composed of multiple CRX sites and one NRL site were more active than equal length CREs composed of only CRX sites (**Fig. 1B**) or only NRL sites (**Fig. S1**). We found that genomic CRX-bound sequences followed a similar pattern [13]. Thus, our previous experiments with systematically varied synthetic CREs show that a sequence context composed of only two types of TFBSs strongly modulates the effects of CRX binding sites. However, it is unclear what kinds of interactions among CRX and NRL sites could account for such context-dependent activity.

**Figure 1.**
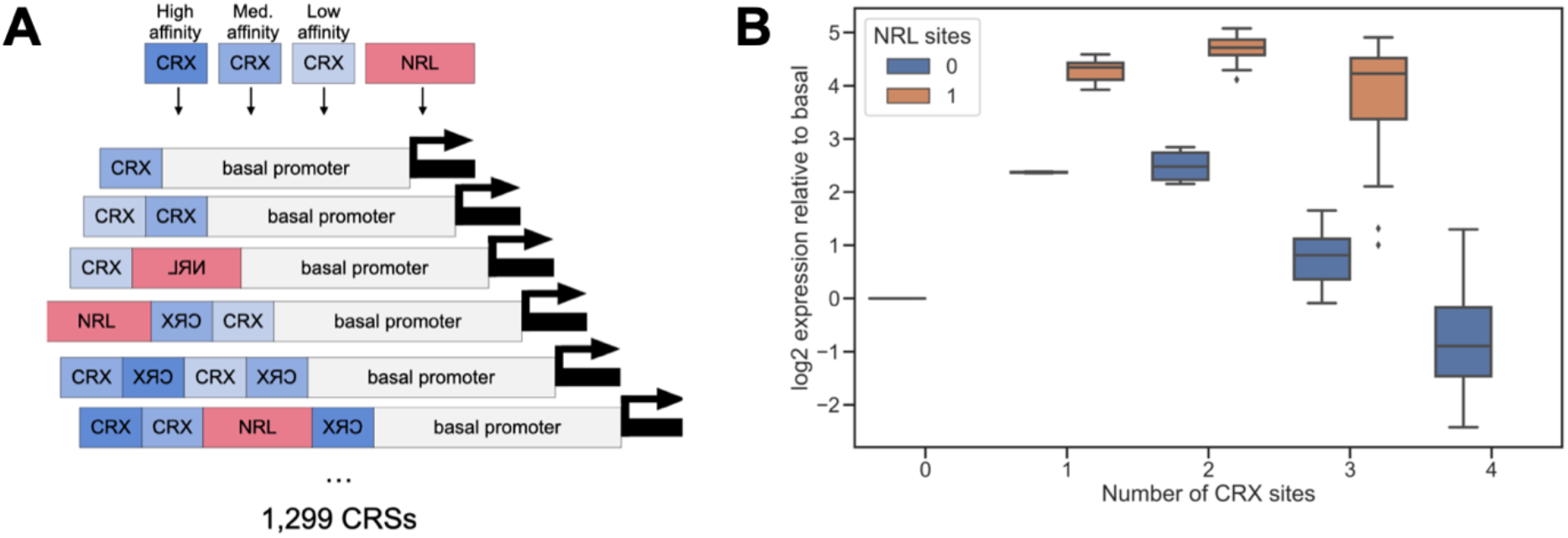
Synthetic CREs sites reveal context-dependent effects of CRX and NRL sites. (A) Design of synthetic CRE MPRA library reported in [13]. Combinations of CRX and NRL sites (up to four TFBSs) were cloned adjacent to either a Rho or a Hsp68 basal promoter. TFBSs could be in either forward or reverse orientation. (B) MPRA activity (y-axis) of CREs composed only of high affinity CRX sites (blue) is consistently lower than that of CREs with high affinity CRX sites and one NRL site (orange), relative to the Rho basal promoter. Sequential addition of high affinity CRX first activates, then represses the Rho basal promoter. Plot shows a subset of the data reported in [13].

To discover interactions among CRX and NRL sites that might explain context-dependent activity, we trained a model using MAVE-NN, a recently published neural network framework that is designed specifically to model data obtained from massively parallel functional assays [47]. A key strength of MAVE-NN is that it deconvolves sequence-function relationships from the confounding effects of experimental non-linearities and noise. Using MAVE-NN, we modeled our synthetic CRE data. As noted above, the synthetic CREs were composed of systematically varied combinations of CRX and NRL sites [13]. Sequences included up to four sites in either the forward or reverse orientation. TFBSs included high, medium, and low affinity versions of CRX sites and the consensus site for NRL (**Fig. 1A**). CREs were cloned upstream of either the murine *Rho* or *Hsp68* basal promoter. The library included all 584 possible combinations of one, two, and three TFBSs, and 715 sequences randomly sampled from all possible combinations of four sites.

We trained MAVE-NN models with different architectures to predict MPRA activities from sequence alone. We reasoned that due to the small number of TFBSs included in the synthetic CREs and the uniform spacing between them, additive models with or without interaction terms would capture most of the effects of CRX and NRL sites on reporter activity. Each model infers a relationship between a reporter gene sequence and its *latent phenotype*, which represents the intrinsic activity of the CRE that is indirectly read out by the MPRA. MAVE-NN simultaneously models (1) the relationship between DNA sequence and the latent phenotype, and (2) the nonlinear relationship between the latent phenotype and the noisy MPRA measurement. MAVE-NN quantifies the performance of the models using an information theoretic measure called *predictive information* [47,48]. Predictive information is the mutual information between the inferred latent phenotype and the MPRA measurement, and it represents how well the model captures the relationship between a reporter gene’s inferred intrinsic activity and its MPRA output. We used predictive information to compare the performance of four different model architectures: (1) an additive model lacking interactions between TFBSs, (2) a nearest-neighbor model that only allows interactions between neighboring TFBSs, (3) a pairwise interaction model allowing interactions between all pairs of TFBSs regardless of spacing, and (4) a ‘black box’ multilayer perceptron model that makes no prior assumptions about the interactions between TFBSs. To train the models, the data was randomly split among training (80%), validation (10%), and test (10%) sets. All performance metrics were computed from the test set. Model parameters for analysis were taken from the best performing model out of multiple random initializations.

Of the three model architectures that included additive and interaction terms, we found that the pairwise interaction model achieved the best overall performance (**Figs. 2A and B**). This model provided 1.8 bits of predictive information, roughly equivalent to an accurate three-way classification of CREs by activity. The predictive information of the pairwise model (1.82 bits) is approximately half that of the multilayer perceptron “black box” model (3.00 bits). The disparity in predictive information between the pairwise and black box models suggests that additional higher order interactions between TFBSs likely account for much of the unexplained activity of the synthetic CREs. However, this unexplained activity likely consists of small discrepancies between sequences with similar activities, because the pairwise model captured a substantial fraction of the variation in reporter activity (**Figs. 2B and C**, R^2^ = 0.889). To understand how additive and interaction effects of TFBSs might explain the context-dependent activity of CRX sites, we examined the parameters of the pairwise model.

**Figure 2.**
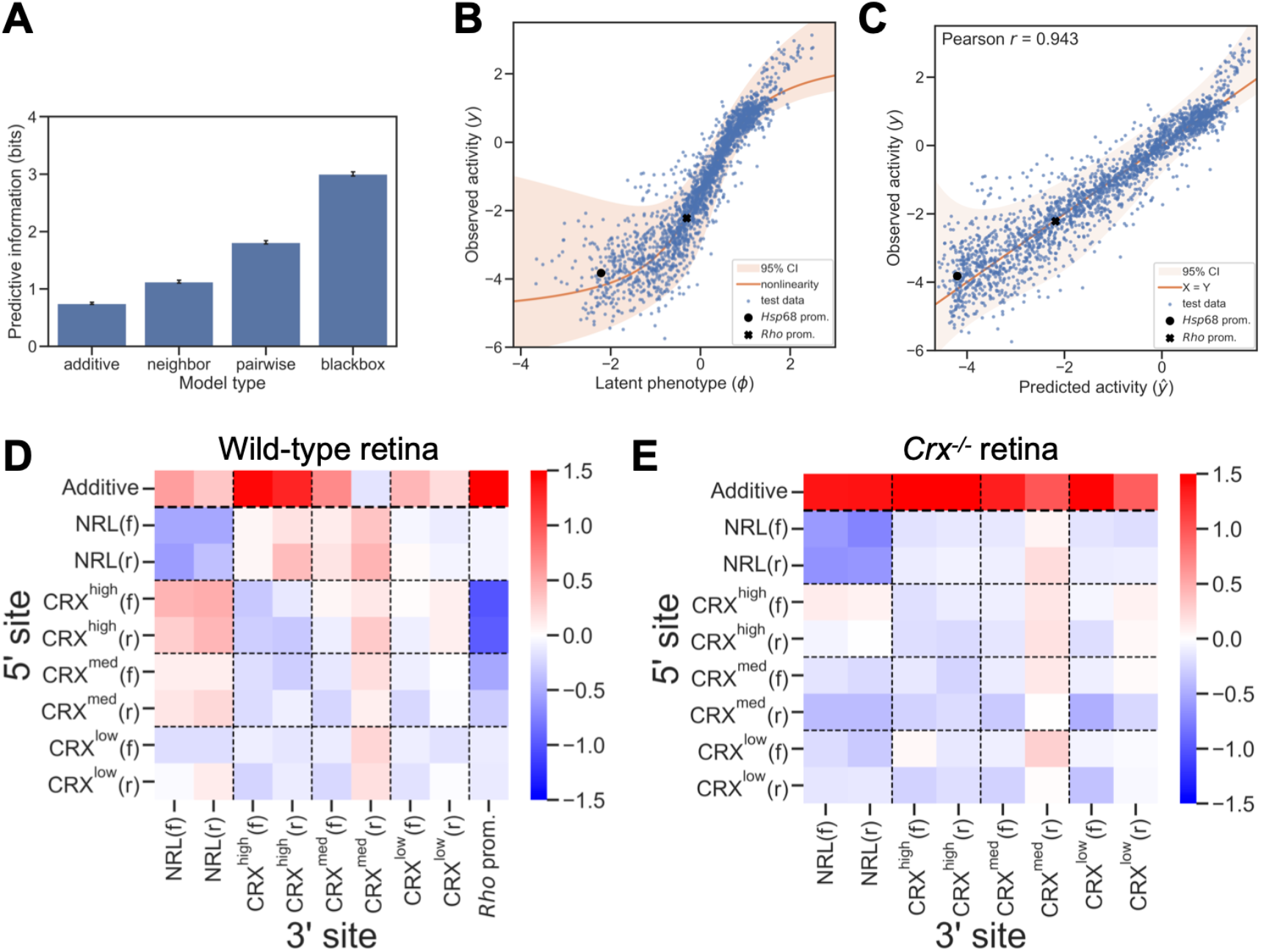
A model of CRX and NRL-driven *cis*-regulatory activity in wild-type retina. (A) The performance of different model architectures (measured as predictive information) fit to MPRA measurements of the CRX-NRL library in wild-type retina. Error bars indicate standard error. (B) The observed activity (y-axis) of test set sequences in wild-type retina compared to the latent phenotype (x-axis) inferred by the pairwise model. (C) The observed activity (y-axis) of test set sequences in wild-type retina compared to the activity predicted by the pairwise model (x-axis). (D) Model parameters for additive and pairwise contributions of CRX and NRL sites and the Rho promoter to activity in wild-type retina, averaged across the four positions in synthetic CREs. For pairwise interactions rows indicate the 5’ site and columns indicate the 3’ site. Forward and reverse orientation of the TFBS is indicated by (f) and (r). (E) Model parameters for additive and pairwise contributions of CRX and NRL sites to activity in Crx^-/-^ retina, averaged across positions and spacings.

We found that the additive contributions of all TFBSs, averaged over all four possible positions in the synthetic CREs, were positive (**Fig. 2D**). This is consistent with the roles of CRX and NRL as transcriptional activators. The average additive contribution of CRX sites increases with site affinity, particularly for sites in the forward orientation. High affinity CRX sites have a stronger positive, additive effect than NRL sites, suggesting CRX is a stronger activator (**Fig. 2D**). The additive terms of the model reflect the expected effects of simple transcriptional activators whose probability of binding to a CRE is determined by the number and affinity of binding sites. However, these positive, additive terms do not account for the context-dependent effects of CRX binding sites observed in the data shown in **Figs. 1B and S1**.

Examining the interaction terms of the model, we observed a pattern of positive, heterotypic cooperativity between CRX and NRL sites, and negative homotypic interactions between binding sites for the same TF (**Fig. 2D**). Negative homotypic interactions are strongest between NRL sites and between high affinity CRX sites, and they decrease with CRX site affinity. CRX sites also show a strong, affinity-dependent negative interaction with the *Rho* basal promoter, which contains three CRX sites [30,49]. Negative homotypic interactions were especially strong between adjacent sites but can be observed at all distances in the synthetic CREs (**Fig. S2D**). Positive interactions between CRX and NRL sites also occur at all distances and depend on binding site affinity. The modeling results suggest that activating and inhibiting interactions between CRX and NRL are the primary determinants of the activity of CREs with binding sites for these two TFs. Because these interactions occur across all distances and not just at neighboring sites, the effects are likely due to multivalent interactions that involve co-factors and not only direct protein-protein interactions between CRX and NRL.

Taken together, the parameters of the pairwise interaction model reveal a *cis*-regulatory grammar that accounts for the observed context-dependent activity of CRX and NRL sites. Consistent with the known roles of CRX and NRL as transcriptional activators, sites for these TFs consistently make positive, independent contributions to activity. However, negative homotypic interactions reduce activation or lead to repression when multiple sites for the same TF are placed together. The repressive effect of negative homotypic interactions can be overcome by the strong positive heterotypic interactions between CRX and NRL. An important feature of this *cis*-regulatory grammar is that additive effects and interactions scale differently with the number of TFBSs. The independent contributions of TFBSs increases linearly with the number of TFBSs, while the interaction effects increase with the square of the number of TFBSs. These differences in scaling have a strong impact on CREs with multiple sites and explain why the replacement of a single binding site can convert a silencer to an enhancer (**Fig. 1B**).

### Positive heterotypic interactions require CRX protein

We previously reported that many genomic and synthetic CREs with CRX binding sites either retain or gain activity in *Crx*^*-/-*^ retina, despite the loss of CRX protein [13]. Activity in *Crx*^*-/-*^ retina still requires intact CRX sites, indicating that another TF, likely the CRX ortholog OTX2, acts at these sites when CRX is absent. To examine how additive and pairwise interactions among TFBSs change in the absence of CRX, we trained a pairwise interaction model on prior data from the synthetic CRE library tested in *Crx*^*-/-*^ retina, with the *Rho* basal promoter. The *Crx*^*-/-*^ model performed similarly to that trained on data from wild-type retina (2.21 bits of predictive information, **Fig. S2 A-C**, R^2^ = 0.900 for predicted versus observed activity). In this model, additive effects of all TFBSs remained positive (**Fig. 2E**), indicating that these CREs continue to be bound by transcriptional activators in *Crx*^*-/-*^ retina. Unlike the model for wild-type retina, the additive contributions of CRX sites did not show a strong dependence on affinity. Negative homotypic interactions remain in *Crx*^*-/-*^ retina, though they are attenuated for CRX sites. Notably, the positive interaction between CRX and NRL sites was absent, indicating that the interaction between these two sites depends specifically on CRX and NRL, and that other TFs that bind these sites in *Crx*^*-/-*^ retina do not interact. Despite the loss of positive cooperativity between CRX and NRL sites, the model suggests that synthetic CREs in *Crx*^*-/-*^ retina maintain or increase their activity due to stronger additive contributions of lower affinity binding sites and a modest attenuation of negative homotypic interactions between CRX sites. The negative homotypic interactions with the *Rho* basal promoter are likely attenuated as well, though we were unable to explicitly model the effects of the *Rho* promoter because there was no data from a library without the *Rho* promoter. Taken as a whole, the model suggests that cooperative interactions depend on the specific identities of the TFs involved, while the positive additive and negative homotypic effects hold more generally among TFs, though with varying effect sizes.

### Additional retinal TFs contribute independently to CRE activity

CRX and NRL are critical for establishing rod photoreceptor identity, and together they drive high expression of a number of key rod photoreceptor genes [34,38,44,46,50]. However, a cooperative interaction between CRX and NRL is not sufficient to explain the context-dependent effects of CRX sites in enhancers, because most CRX-bound enhancers do not contain a copy of the NRL motif [18,42]. To investigate how other TFBSs contribute to the activity of CREs that contain CRX sites, we designed a new library of 6,600 synthetic CREs. The library included TFBSs for CRX, NRL, as well as sites from three other photoreceptor-enriched motif families: NEUROD1, RORB, and motifs representing SP4 or MAZ [41,51–53]. We previously found that motifs for these TFs were enriched in CRX-bound enhancers, but not silencers [18]. We sought to discover whether these additional TFBSs interact cooperatively with CRX sites, or whether they contributed independently to enhancer activity. We designed synthetic CREs composed of five sites and we systematically varied the TFBSs composition across the library. Each CRE included either two or three CRX sites and one or two sites for two additional TFs (**Fig. 3A**). Synthetic CREs were cloned upstream of the *Rho* minimal promoter and tested by MPRA with three replicate transfections in explanted retinas (mean R^2^ between replicates = 0.950, **Fig. S3A**). We used the data to train different models and again found that the pairwise interaction model performed better than the additive or nearest-neighbor models (predictive information = 2.65). No additional performance was gained from the black box model (**Fig. S3B**). The pairwise model captured most of the variance in CRE activity (**Fig. 3B and C**, R^2^ between predicted and observed expression = 0.970).

**Figure 3.**
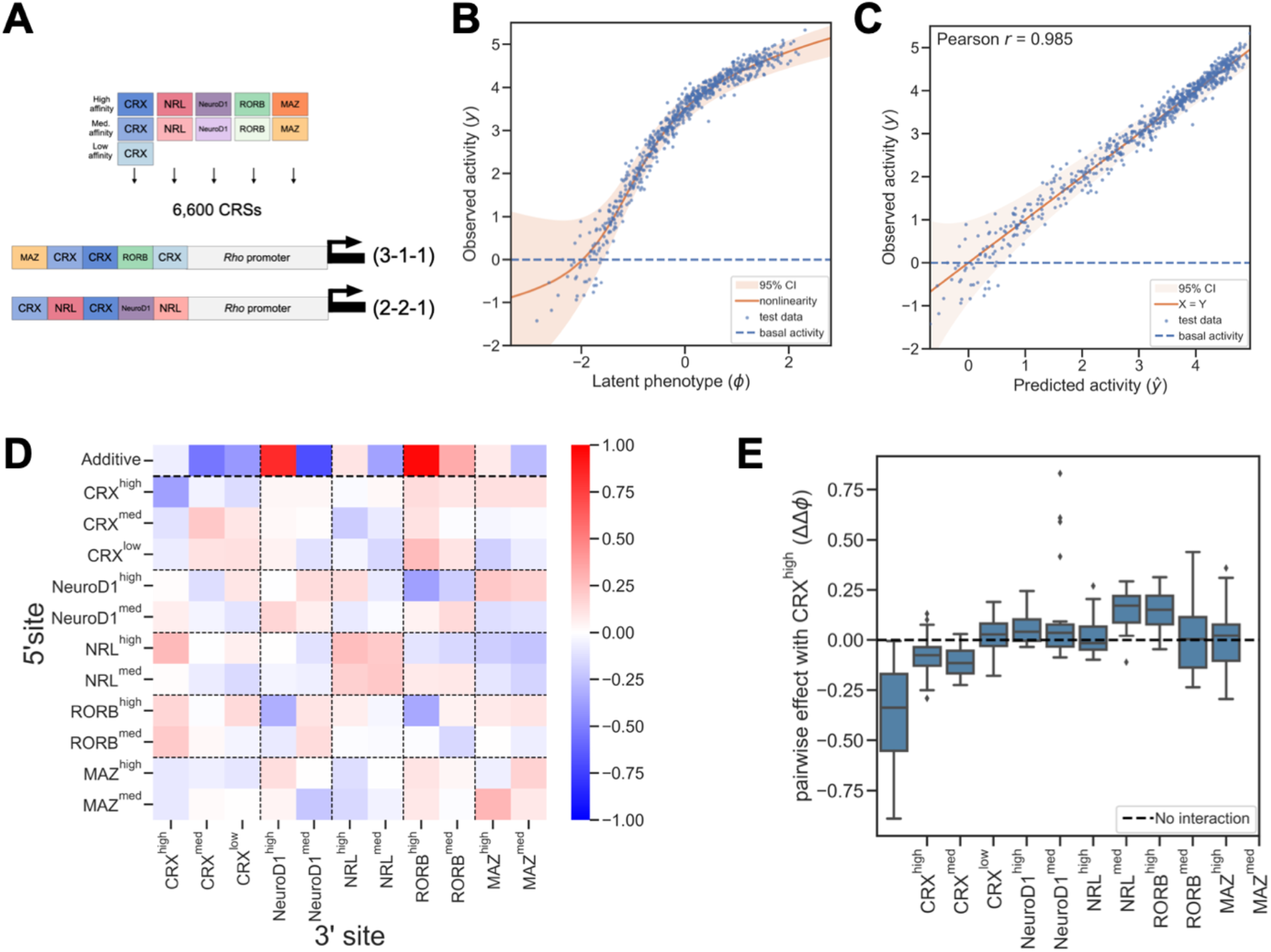
A model of *cis*-regulatory activity driven by diverse TFBSs in wild-type mouse retina. (A) Design of MPRA library of synthetic CREs with additional lineage-specific TFBSs. CREs contained five sites placed adjacent to the Rho basal promoter. Each CRE contained either three CRX sites and two sites for other TFs (3-1-1) or two CRX sites, two sites for another TF, and one site for a third TF (2-2-1). (B) Observed activity (y-axis) of test set sequences compared to the latent phenotype (x-axis) predicted by the pairwise model. (C) Observed activity (y-axis) of test set sequences compared to the activity predicted by the pairwise model (x-axis). (D) Model parameters representing additive and pairwise contributions of TFBSs averaged across positions. (E) Distributions of interactions with high-affinity CRX binding sites across all positions, broken down by partner TF.

Examining the average additive effects of TFBSs in the pairwise interaction model, we found that higher affinity sites for NRL, NEUROD1, RORB, and MAZ contributed positively to activity, while lower affinity sites had weaker positive effects or negative effects on activity (**Fig. 3D**). High affinity sites for NEUROD1 and RORB made especially strong additive contributions to activity. In contrast to the models above, the additive contributions of CRX sites in this model were negative. This is likely due to the design of the library, which only includes CREs with two or three CRX sites, making it difficult to deconvolve additive effects of individual CRX sites from the effects of negative homotypic interactions between CRX sites. The interaction terms of the model exhibit a mixture of moderate, positive and negative effects that depend on the affinity and order of the TFBSs (**Figs. 3D and S3C**). We looked specifically for interactions between other TFs and CRX by examining the interaction terms between high affinity CRX sites and each type of TFBS (**Fig. 3E**). As with the prior model, there were strong, affinity-dependent negative homotypic interactions between CRX sites. High affinity NRL sites interacted positively with CRX sites only in some positions, in contrast to what we observed in the CRX + NRL library. This could be due to differences in CRE structures between the two libraries or to differences in the NRL motif sequence (see Methods). Sites for RORB showed consistent positive interactions with CRX sites across positions, while sites for NEUROD1 and MAZ generally did not interact with CRX sites. The results of this model suggest that, while CRX does cooperatively interact with some TFs, such interactions are not necessary to overcome the repressive effects of negative homotypic interactions. Instead, the additive contributions from diverse TFBSs can shift the balance towards activation.

### Balance between positive and negative interactions can explain context-dependent effects of binding sites for transcriptional activators

The MAVE-NN models suggest that the context dependency of sites for transcriptional activators like CRX and NRL can be explained by the balance between the negative effects of homotypic interactions and the positive effects of individual TFBSs and heterotypic interactions between them. These effects create context dependence without the need for repressor TFBSs or dual-function TFs with distinct activation and repression domains. Instead, the results presented above suggest that some transcriptional activators self-inhibit when present at higher occupancy on a CRE. The negative effects of self-inhibition can be overcome by positive cooperativity with a different TF, or by the non-cooperative action of a diverse collection of TFs that, as a collective, engage in fewer negative homotypic interactions. Under this model, the TFBS composition at enhancers and silencers shifts the balance between these effects in favor of either activation or repression. At enhancers, positive cooperativity and the independent contributions of diverse activator TFBSs outweigh the effects of negative homotypic interactions, while at silencers negative homotypic interactions predominate.

To demonstrate how context dependence is achieved under such *cis*-regulatory grammar of balanced effects, we implemented a simplified model that expresses CRE activity as the sum between positive and negative contributions of activator TFBSs. This model recapitulates the non-monotonic relationship we observed between the number CRX sites and CRE activity (**Fig. 1B**). In the model, we assume that CRE activity is the sum of (1) positive, additive contributions from sites for transcriptional activators, (2) positive cooperativity between sites for different TFs, and (3) negative interactions between sites for the same TF. For CREs composed only of sites for two different TFs, as in Fig. 1, this sum is

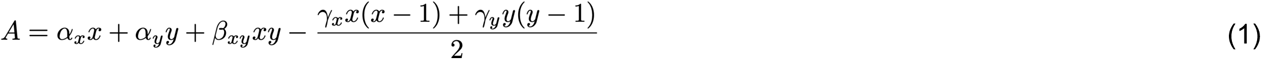

where *A* is activity of a CRE, *x* is the number of sites for the first TF, *y* is the number of sites for the second TF, and *α, β, γ* are weights reflecting the relative strength of each contribution to activity. The first two terms represent the additive contribution of each TFBS, the third term represents positive cooperativity between all pairs of sites for different TFBSs, and the final term represents negative interactions between all pairs of sites for the same TF.

We calculated the expected activities of all possible CREs with up to four sites (**Fig. 4A**), making the simplifying assumption that the relative strengths of the different terms in eq. 1 are similar and setting all weights equal to 1. The simulated activities recapitulate patterns of expression observed in the library of synthetic CREs with combinations of CRX and NRL sites. Starting with the basal promoter alone (indicated by zero), increasing the number of sites for a single TF leads first to an increase and then a decrease in activity (leftmost column or top row in **Fig. 4A**, compare with **Figs. 1B** and **S1**). The highest activities are obtained from CREs with combinations of sites from both TFs. In the model, a CRE with four CRX sites is repressive. Replacing one of those sites with an NRL site converts the CRE from a silencer to an enhancer, an effect also observed in the data (**Figs. 4A** and **1B**.) While our model relies on simplifying assumptions that are unlikely to fully hold *in vivo*, it successfully recapitulates the major trends observed in our data.

**Figure 4.**
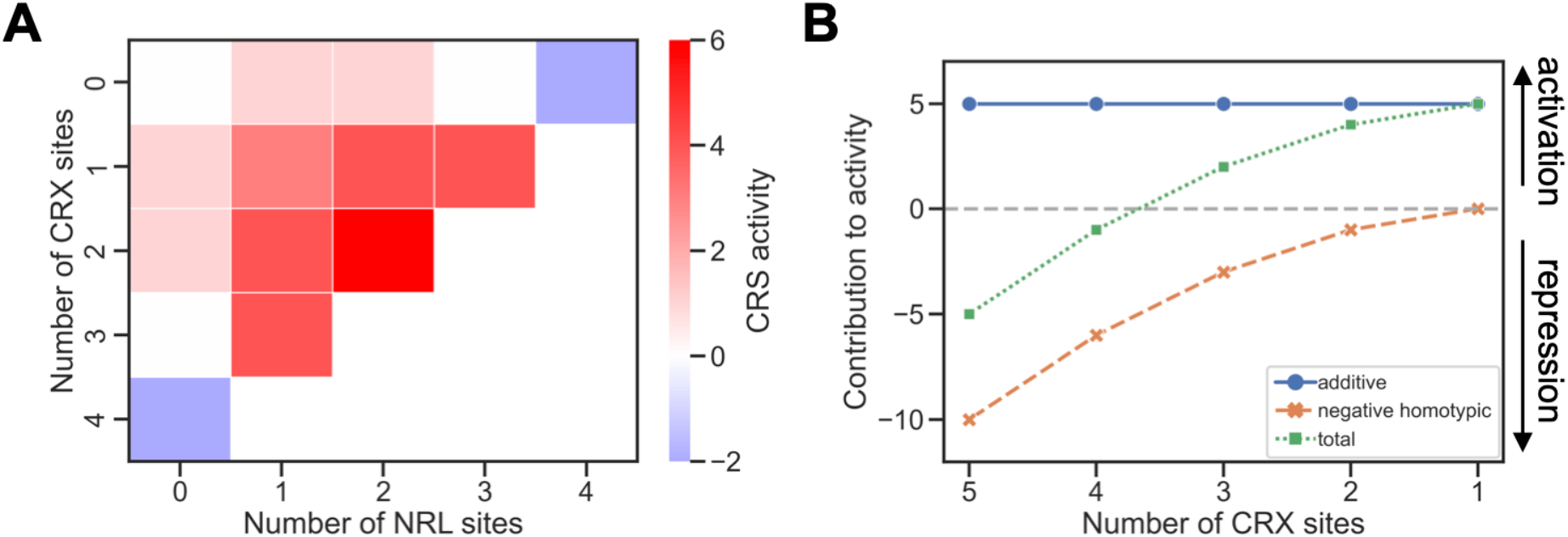
Simplified balance model of context-dependent effects of binding sites for transcriptional activators. (A) Simulated CRE activities calculated by eq. 1 for sequences with up to four TFBSs for CRX or NRL. Stepwise addition of sites for a single TF first increase then decrease activity. The first two columns show predicted expression of CREs with only CRX sites or with CRX sites plus one NRL site. Compare with the measured values in Fig. 1B. (B) Model of CREs with five TFBSs shows how TF diversity reduces negative homotypic interactions and increases CRE activity. As CRX sites are replaced with sites for different TFs, TF diversity increases (x-axis) and the number of negative homotypic interactions decreases (orange crosses) and the overall CRE activity increases (blue squares). The total additive contribution of TFBSs (green circles) is equal to the total number of TFBSs and remains constant.

We also modeled the effects of TFBS diversity in the absence of cooperative interactions. The MAVE-NN model of synthetic CREs with five TFBSs suggests that a diversity of TFBSs can shift the balance in favor of activation, even without cooperative interactions (**Fig. 3D**). For a CRE composed of a given number of sites, greater TFBS diversity reduces negative homotypic interactions. As a result, the independent positive effects of each TFBS predominate. To model these effects, we assumed that (1) bound TF activators always make positive, independent contributions to activity and (2) all TFs engage in negative homotypic interactions. We calculated the sums of positive and negative effects for CREs with five total TFBSs, but different numbers of CRX sites (and therefore varying amounts of negative homotypic interactions). In these simulated CREs, the total additive contribution is constant and equal to the total number of TFBSs (**Fig. 4B**, blue circles). As the diversity of the TFBSs increases, the number of negative homotypic interactions is reduced (**Fig. 4B**, orange crosses). A CRE with five CRX sites is therefore highly repressive, while replacing some CRX sites with different TFBSs increases the activity of the CRE (**Fig. 4B**, green squares). This simplified model demonstrates how strong activity can be achieved by the independent effects of diverse TFBSs, even in the absence of cooperative interactions. This model suggests an explanation of our prior observation that CRX-bound strong enhancers have more diverse TFBSs than CRX-bound silencers [18].

## Discussion

Because the effects of a TFBS often strongly depend on local sequence, the activity of *cis*-regulatory DNA is not a simple function of TFBS composition. To accurately predict the activities of *cis*-regulatory sequences and the effects of genetic variants that occur within them, we need models of *cis*-regulatory grammar that accurately account for the influence of sequence context. Different occurrences of a TF binding motif can differ in their effects due to post-translational modifications of TFs [54,55], the presence of other co-bound factors [5,6,9,11,12,19,28,56–59], and binding by different TFs with similar sequence specificities [10,60,61]. Our results suggest that context dependence can also be determined by the overall balance between independent and interaction effects of individual sites for transcriptional activators. At the core of this model is a distinction between additive, independent effects of individual TF molecules and effects of interactions between molecules. In the case of CRX, the independent and interaction effects influence *cis*-regulatory activity in opposite directions, with CRX molecules independently contributing to activation while engaging in repressive homotypic interactions with one another. The independent, activating effects scale linearly with the number of binding sites, while the number of repressive homotypic interactions scales with the square of binding site number. As the number of binding sites in a CRE increases, negative homotypic interactions grow faster than the activating effects of individual binding sites. As a result, sequences with many CRX sites are likely to act as silencers, a pattern that we observe with both synthetic CREs and genomic CRX-bound sequences [13,18].

In this model, other TFs can influence whether a CRX-bound CRE will activate or repress transcription in two ways. First, a TF like NRL may form positive cooperative interactions with CRX. Positive cooperative interactions also scale with the square of the numbers of binding sites, thus the addition of relatively few sites for a cooperating TF can shift the balance towards activation. This explains our observation that, for a repressive CRE with four CRX sites, converting one position to an NRL site changes activity of the CRE from repression to activation (**Fig. 1B**). This effect in synthetic CREs is consistent with our previous finding that genes near CRX-bound loci that are co-bound by NRL are more highly expressed than genes near regions bound by CRX alone [13]. In the second way, the presence of other bound TFs at a CRE can also overcome negative homotypic effects by making strong additive contributions, without cooperativity, as in the case of NEUROD1. At a CRE, having fewer sites for a single TF and more sites for a variety of TFs minimizes negative homotypic interactions and allows the additive effects of TFBSs to predominate. The repressive effects of homotypic interactions can thus be overcome by positive cooperativity, or by the presence of diverse TFBSs.

This model suggests an explanation for our prior observation that CRX-bound enhancers often have more diverse TFBSs than CRX-bound silencers [18]. Genomic sequences that act as silencers tend to contain clusters of multiple CRX sites and few sites for other TFs, while genomic enhancers tend to contain sites for a variety of photoreceptor TFs [13,18,42]. These patterns are recapitulated in synthetic CREs whose TFBS composition is systematically varied. We previously quantified the role of TFBS diversity in distinguishing silencers from enhancers using a phenomenological metric that we called information content, which considers the number and diversity of TFBSs in a CRE [18]. High-information-content sequences have a greater number and diversity of TFBSs and are more likely to be enhancers, while silencers often have low information content because they lack TFBS diversity. Our MAVE-NN models suggest that high-information-content sequences are enhancers because binding site diversity allows the additive contributions of TFBSs to outweigh negative homotypic interactions. This contrasts with low-diversity silencers, which our model suggests are dominated by negative homotypic interactions. TFBS diversity is a feature of enhancers in other cell types, where a similar balance between positive additive effects and negative homotypic interactions may occur [62].

In our models, negative homotypic interactions strongly influence the context-dependent effects of binding sites for several photoreceptor TFs, particularly CRX and NRL. The existence of such interactions is supported by data from both synthetic CREs and genomic sequences. Sequential addition of CRX or NRL binding sites upstream of the *Rho* basal promoter first increases, then decreases transcription, sometimes below basal levels (**Fig. 1B and S1**) [13]. Genomic CRX-bound sequences that act as silencers when measured by MPRA have more copies of the CRX motif than sequences that act as enhancers [13,18,42]. We have shown that this silencing activity depends on both CRX motifs and CRX protein [13]. Similar negative homotypic effects have been reported for several TFs, including liver specific factors [63], yeast Gcn4 [64], pluripotency TFs [16,65], and Sp3 [66]. These findings suggest some TFs may self-inhibit or recruit repressors when they highly occupy certain CREs [20,67,68]. The homeodomain TF WUSCHEL activates transcription as a monomer at low concentration but forms repressive dimers at higher concentration [69,70]. A recent study found that homeodomain TFs are enriched in activation domains that also exhibit the ability to repress [71]. Our model of context dependency suggests that the balance between positive effects and negative homotypic interactions can account for the dual activities of some TFs, without the need to invoke dedicated repressors.

## Methods

### Model fitting

For the CRX + NRL library, binding site arrangements and MPRA activities were extracted from Database S3 of [13]. For the library with CRX, NEUROD1, NRL, RORB, and MAZ sites (CDNRM library), the MPRA experiment was performed as described below. Data files are described below under **Data availability**. To encode arrangements of TFBSs as input sequences for MAVE-NN, we used single letters to represent each type of binding site. To create input sequences of uniform length for the CRX + NRL library, dummy binding sites labeled “O” were prepended to each arrangement to render all CREs four sites long, and an additional letter indicating the basal promoter (*Rho* or *Hsp68*) was then appended to the end. Models were trained using mave-nn package version 1.01 until convergence on the processed data, using hyperparameters given in **Table 1**. The models were specified using the Skewed-T GE noise model with a heteroskedasticity order of 2. We used the consensus gauge with basal *Hsp68* as the consensus sequence to obtain parameters from the models trained on CRX-NRL data in wild-type retina, and the uniform gauge for the remaining models. To ensure consistent training outcomes, we trained each model from a large number of random initializations (25 for the CRX + NRL library in wild-type retina, 20 for the library in *Crx*^*–/-*^ retina, and 50 for the CDNRM library), with the numbers chosen to achieve maximum performance and reproducibility. We picked the best-performing model of each type for further evaluation. Model performance was evaluated by cross-validation with an 80-10-10 percent training-validation-test set split. The measurements were split randomly between sets and the same split was used for all random initializations.

**Table 1:**
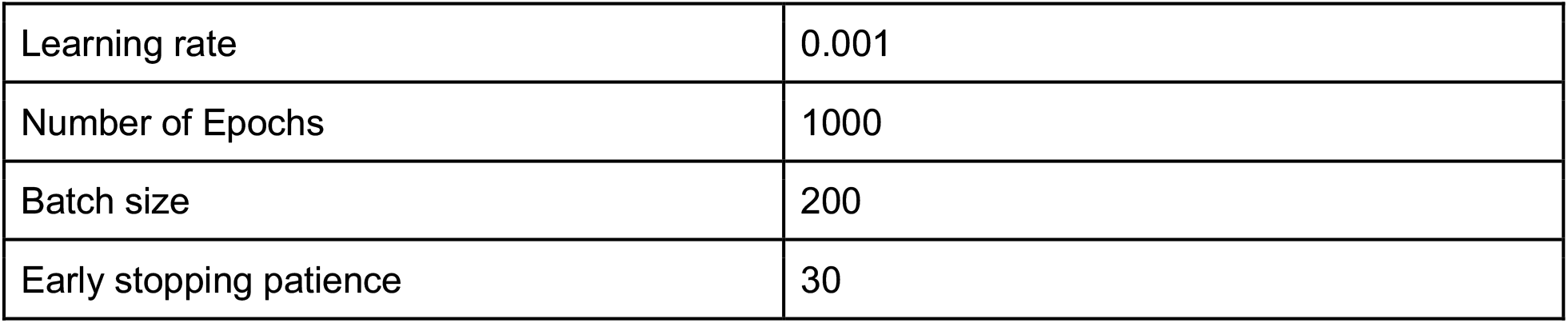
Hyperparameters used to fit models to building block data

|  |  |
| --- | --- |
| Learning rate | 0.001 |
| Number of Epochs | 1000 |
| Batch size | 200 |
| Early stopping patience | 30 |

**Table 2:** Hyperparameters used to fit models to *Rho* promoter mutagenesis data

|  |  |
| --- | --- |
| Learning rate | 0.0005 |
| Number of Epochs | 2000 |
| Batch size | 100 |
| Early stopping patience | 25 |

### CDNRM library design

We designed a library of 6600 synthetic CREs composed of combinations of binding sites for CRX, NRL, NEUROD1, RORB, and MAZ. The library was designed to vary TFBS diversity around CRX sites. It contained all possible arrangements of either 3 sites for CRX and 1 site each for two other TFs (3-1-1 sequences); or 2 sites for CRX, 2 sites for a second TF, and 1 site for a third TF (2-2-1 sequences). CRX sites in a CRE were either all high affinity or a mixture of affinities. TFBS orientation was held constant. High, medium and low affinity CRX sites were those used in the CRX + NRL library [13]. The NRL, NEUROD1, RORB, and MAZ motifs were randomly selected from sites identified in genomic strong enhancers that lose activity when the corresponding motif is deleted [18]. Note that NRL sites in the CDNRM library vary from the site in the CRX + NRL library, but all sites for the synthetic CREs have been shown to be active in genomic sequences. Binding site sequences were padded to make all motifs 12 bp, then a constant buffer sequence was added (AGCTAC<motif>GT) to create a 20 bp “building block” that maintains helical spacing when sites were combined, similar to our procedure for prior libraries of synthetic CREs [13,16,65]. The 12 bp motifs used with the core motif underlined, are: high affinity CRX, TG<u>CTAATCCC</u>AC; medium affinity CRX, TG<u>CTAAGCCA</u>AC; low affinity CRX, TG<u>CTGATTCA</u>AC; high affinity NRL: <u>AATTTGCTGAC</u>C; med NRL, <u>GGCCTGCTGAC</u>C; high affinity NEUROD1, C<u>AACAGATGGT</u>A; medium affinity NeuroD1, C<u>GGCAGGTGGT</u>A; high affinity RORB, <u>AATTAGGTCAC</u>T; medium affinity RORB, <u>ATCTGGGTCAG</u>T; high affinity MAZ, <u>GGGGGAGGGGG</u>G; medium affinity MAZ, <u>GCGGGCGGGGG</u>G.

### MPRA library cloning

Synthetic CREs were each represented in the library with 3 unique barcodes. As standards, the library included 20 genomic sequences taken from [18] that span the dynamic range of the MPRA and 150 scrambled sequences as negative controls. The *Rho* basal promoter was tagged with 90 barcodes to ensure precise measurement of basal levels. Barcoded CREs were synthesized as two sub-libraries on a single chip using custom oligonucleotide synthesis from Agilent Technologies. The oligonucleotide libraries were cloned as previously described [18]. Briefly, we amplified oligos using either primer pairs MO563 (GTAGCGTCTGTCCGTGAATT) and MO564 (CTGTAGTAGTAGTTGGCGGC) or RZFP3 (TCTAGACTGCGGCTCGAATT) and RZFP4 (AGATCTAATGCATACGCGGC), and cloned them into the vector pJK03 (AddGene #173,490). The rod-specific *Rho* promoter, the *DsRed* reporter gene, and a multiplexing barcode (mBC) was cloned between the synthetic sequence and the cBC. One sub-library was assigned mBC TAGTAACGG, the other was assigned CCTACTAGT. The final plasmid libraries were mixed together at equimolar concentrations.

### Retinal explant electroporation

Animal procedures were performed in accordance with a Washington University in St Louis Institutional Animal Care and Use Committee-approved vertebrate animals protocol. Electroporations into retinal explants into P0 CD-1 mice and RNA extractions were performed as described previously [13,18,42,49]. We performed three replicate electroporations. cDNA and the input plasmid pool was sequenced on the Illumina NextSeq platform. We obtained an average sequencing depth of >675 reads per barcode.

### MPRA data processing

Sequencing reads were filtered to retain only perfect barcode matches. After filtering we retained 95% of sequencing reads. Barcodes with fewer than 50 reads in the plasmid pool were considered missing and removed. Barcode read counts were normalized by total sample reads. MPRA activity scores for each replicate were calculated by dividing RNA by DNA values, averaging across barcodes for each CRE, then normalizing to the activity of the basal promoter [18]. Replicates were averaged and the log_2_ transformed values were used for model training.

## Data availability

Processed data files and code are available at https://github.com/mawsterlab/synthetic_promoter_mavenn. FASTQ files for the MPRA experiments with the CDNRM library are deposited with the NCBI Gene Expression Omnibus (GEO) under accession number GSE225867.

## Funding

This work was supported by National Institutes of Health grants R01 GM121755 to M.A.W.; R01 GM092910 to B.A.C.; EY030075, HL149961, and MH122451, to J.C.C.; and F31HG011431 to R.Z.F. B.A.C. is on the scientific advisory board of Patch Biosciences.

## Supplemental Figures

**Figure S1:**
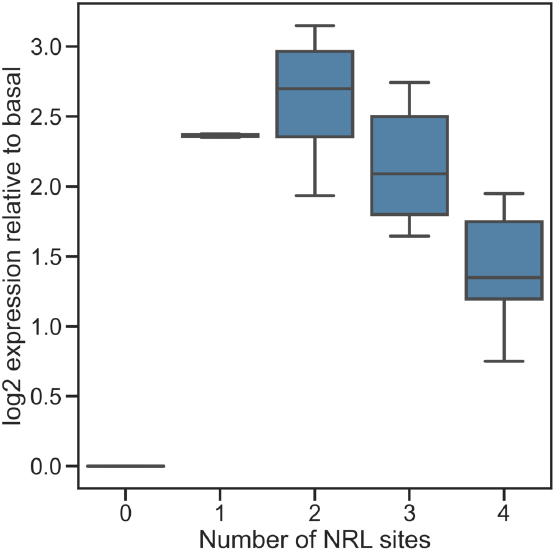
Increasing NRL sites reduces MPRA activity in synthetic CREs. Synthetic CREs composed only of NRL sites show an increase, then a decrease in activity relative to the *Rho* basal promoter as the number of sites is increased. Plot shows a subset of the data reported in [13].

**Fig. S2.**
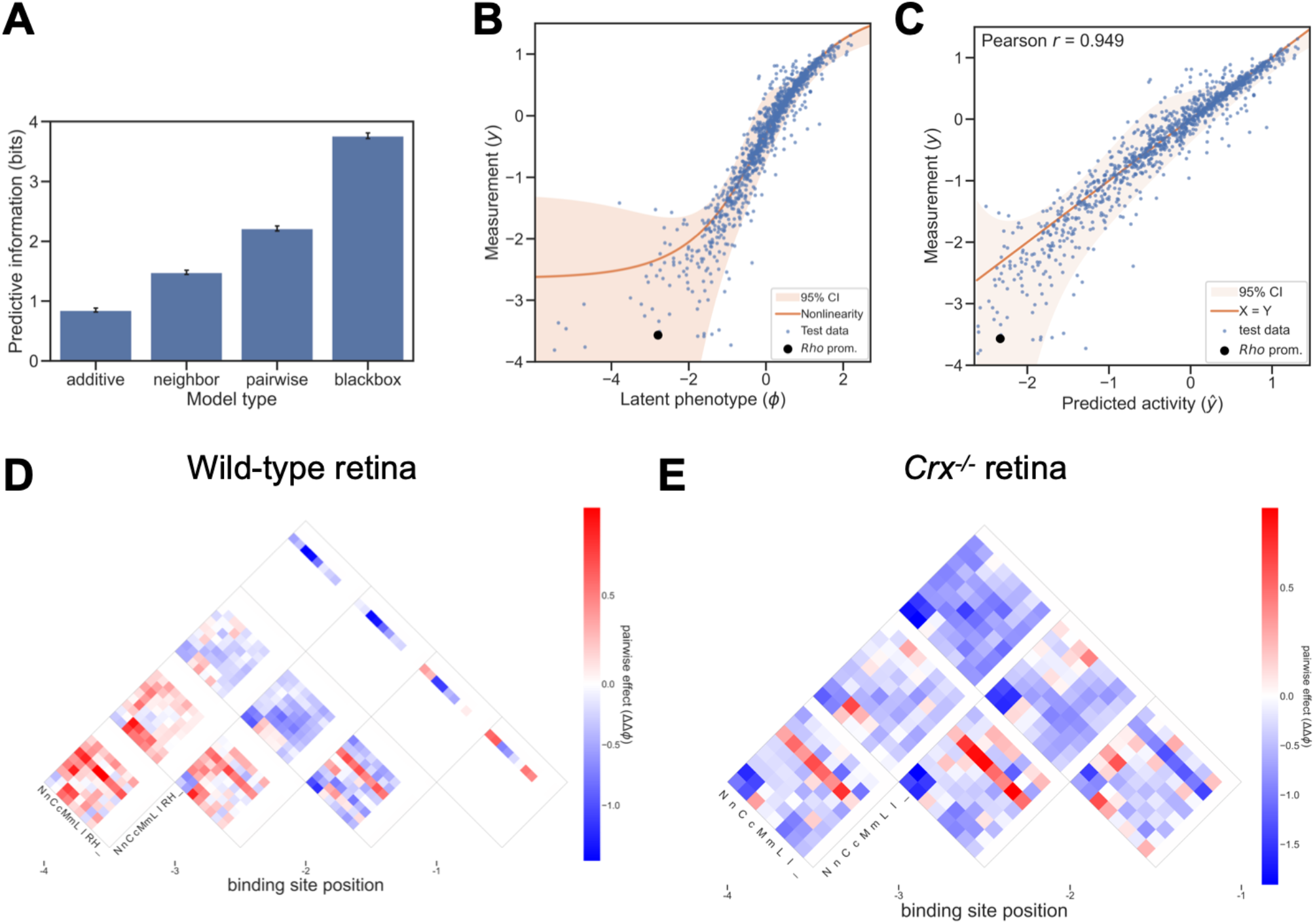
A model of CRX and NRL-driven *cis*-regulatory activity in *Crx*^*-/-*^ retina. (A) The performance of different model architectures (predictive information) fit to MPRA measurements of synthetic CREs in *Crx*^*-/-*^ retina. (B) Observed activity (y-axis) of test set sequences in *Crx*^*-/-*^ retina compared to the latent phenotype (x-axis) predicted by the pairwise model. (C) Observed activity (y-axis) of test set sequences in *Crx*^*-/-*^ retina compared to the activity predicted by the pairwise model (x-axis). (D) Model parameters for position-specific pairwise contributions of CRX and NRL sites in wild-type retina. Forward and reverse orientation of binding sites is indicated by capital or lower case letter. CRX sites are either high (C or c), medium (M or m), or low (L or l) affinity. NRL sites are labeled N or n and the *Rho* promoter is labeled R. There are no model parameters for *Hsp68* (H, used as the overall basal sequence) or the placeholder site _ used to equalize the lengths of input sequences. See methods for details. (E) Position-specific pairwise contributions of CRX and NRL sites to activity in *Crx*^*-/-*^ retina.

**Fig. S3.**
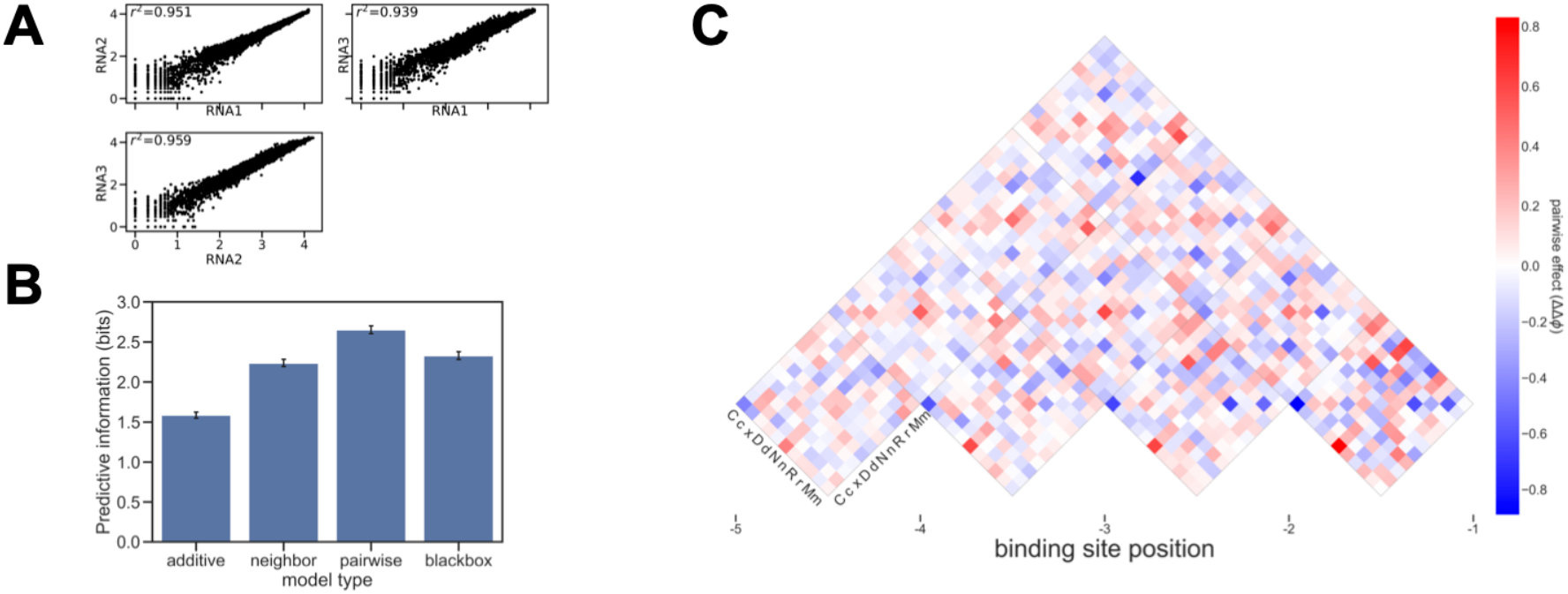
A model of *cis*-regulatory activity driven by diverse TF binding sites in wild-type retina. (A) Reproducibility of MPRA measurements across three replicates. (B) Performance of modes fit to measurements of the MPRA library of CREs composed of five TFBSs, expressed in terms of predictive information. (C) Position-specific pairwise contributions of diverse TF binding sites. Model parameters shown are the average of multiple training runs from different random starts as described in the methods. Capital and lowercase letters represent high and medium affinity sites for CRX (C), NEUROD1 (D), NRL (N), RORB (R), and MAZ (M). Low affinity CRX sites are represented by x.

## Notes

https://github.com/mawsterlab/synthetic_promoter_mavenn

### Competing Interest Statement

The authors have declared no competing interest.

